# Characteristics and origins of non-functional *Pm21* alleles in *Dasypyrum villosum* and wheat genetic stocks

**DOI:** 10.1101/332841

**Authors:** Shanying Zhu, Huagang He

## Abstract

Most *Dasypyrum villosum* resources are highly resistant to wheat powdery mildew that carries *Pm21* alleles. However, in the previous studies, four *D. villosum* lines (DvSus-1 ∼ DvSus-4) and two wheat-*D. villosum* addition lines (DA6V#1 and DA6V#3) were reported to be susceptible to powdery mildew. In the present study, the characteristics of non-functional *Pm21* alleles in the above resources were analyzed after Sanger sequencing. The results showed that loss-of-functions of *Pm21* alleles *Pm21-NF1* ∼ *Pm21-NF3* isolated from DvSus-1, DvSus-2/DvSus-3 and DvSus-4 were caused by two potential point mutations, a 1-bp deletion and a 1281-bp insertion, respectively. The non-functional *Pm21* alleles in DA6V#1 and DA6V#3 were same to that in DvSus-4 and DvSus-2/DvSus-3, respectively, indicating that the susceptibilities of the two wheat genetic stocks came from their *D. villosum* donors. The origins of non-functional *Pm21* alleles were also investigated in this study. Except the target variants involved, the sequences of *Pm21-NF2* and *Pm21-NF3* were identical to that of *Pm21-F2* and *Pm21-F3* in the resistant *D. villosum* lines DvRes-2 and DvRes-3, derived from the accessions GRA961 and GRA1114, respectively. It was suggested that the non-functional alleles *Pm21-NF2* and *Pm21-NF3* originated from the wild-type alleles *Pm21-F2* and *Pm21-F3*. In summary, this study gives an insight into the sequence characteristics of non-functional *Pm21* alleles and their origins in natural population of *D. villosum*.

## Introduction

*Dasypyrum villosum* Candargy (2*n* = 2*x* = 14, VV) is a diploid species native to the Mediterranean region. *D. villosum* not only has good tiller ability and high grain protein content, but also provides tolerance to drought and cold stresses, and resistance to multiple wheat diseases, such as powdery mildew, rusts, eyespot, Take-All, and wheat spindle streak mosaic disease (De Pace et al. 2011; Wang et al. 2017). Hence, as a wild relative, *D. villosum* has been concerned to be an important resource for improvement of common wheat (*Triticum aestivum* L., 2*n* = 6*x* = 42, AABBDD).

In 1995, a wheat-*D. villosum* translocation line T6AL.6VS (6V#2S) was developed from an amphidiploid of durum wheat and *D. villosum* accession provided by Cambridge Botanical Garden, United Kingdom. The line T6AL.6VS carries the powdery mildew resistance gene *Pm21* that confers highly resistant to all tested races of *Blumeria graminis* f. sp. *tritici* (*Bgt*) (Chen et al. 1995). Since then, more than 20 varieties carrying *Pm21* have been developed and applied. Especially, in the middle and lower reaches of the Yangtze River Valley, the most rampant area of powdery mildew in China, wheat varieties carrying *Pm21* are been planted more widely than the ones carrying *Pm2a* or *Pm4a* that are gradually losing their resistance in this region (Bie et al. 2015; He et al. 2017).

Besides 6V#2S carrying *Pm21*, other wheat genetic stocks involved in chromosome 6V or 6VS of *D. villosum* have been developed by different research groups (Sears 1953; Hyde 1953; Chen et al. 1995; Li et al. 2005; Liu et al. 2011; Lukaszewski and Cowger 2017). Interestingly, among them, the addition lines DA6V#1 (Sears 1953) and DA6V#3 (Lukaszewski, unpublished) were reported to be susceptible to wheat powdery mildew (Qi et al. 1998; Liu et al. 2011). It is difficult to understand why these materials are susceptible since no any susceptible *D. villosum* donor was found in the previous study (Qi et al. 1998).

Recently, four seedling-susceptible *D. villosum* lines (DvSus-1 ∼ DvSus-4) were screened from 110 accessions in our study. Based on the finding of susceptible *D. villosum* resources, a fine genetic map of *Pm21* was constructed, and then *Pm21* was successfully characterized as a typical coiled-coil, nucleotide-binding site, leucine-rich repeat (CC-NBS-LRR) protein-encoding gene (He et al. 2017; He et al. 2018). Thus, one basic question arises: what are the characteristics of non-functional *Pm21* alleles in the susceptible *D. villosum* lines and related wheat genetic stocks. Moreover, given that susceptible *D. villosum* resources are very rare, it is greatly interesting to clarify whether all or some of the susceptible *D. villosum* lines and wheat genetic stocks share the same mutation(s) of *Pm21* alleles and whether we can trace the origins of non-functional *Pm21* alleles in natural population of *D. villosum*. Therefore, in the present study, we made an attempt to answer the questions above via sequencing and analysis of *Pm21* alleles.

## Materials and methods

### Plant materials and growth conditions

The resistant *D. villosum* line DvRes-1 carrying *Pm21* was provided by Prof. Peidu Chen, Cytogenetics Institute of Nanjing Agricultural University (CI-NAU). Other *D. villosum* lines used in this study were provided by Germplasm Resources Information Network (GRIN) and Genebank Information System of the IPK Gatersleben (GBIS-IPK). The wheat-*D. villosum* disomic addition lines DA6V#1 and DA6V#3 were kindly provided by GRIN and Prof. Bernd Friebe, Wheat Genetic and Genomic Resource Center, Kansas State University (WGGRC-KSU), respectively (Table 1). The powdery mildew resistant wheat cultivated variety (cv.) Yangmai 18, a translocation line carrying *Pm21*, and the susceptible cv. Yangmai 9 were both developed in Yangzhou Academy of Agricultural Sciences (YAAS), Yangzhou, Jiangsu province. Plants were grown in greenhouse with LED light under long-day condition (16 h light/8 h dark) at 24°C.

**Table 1.** *D. villosum* lines and wheat genetic stocks used in this study. CI-NAU: Cytogenetics Institute, Nanjing Agricultural University. GBIS-IPK: Genebank Information System of the IPK Gatersleben. GRIN: Germplasm Resources Information Network. WGGRC-KSU: Wheat Genetic and Genomic Resource Center, Kansas State University

| Line | Allele<sup>a</sup> | Powdery<br>mildew<br>response | Origin | Original<br>accession | Provider |
| --- | --- | --- | --- | --- | --- |
| DvRes-1 | *Pm21* | Resistant | United Kingdom | Unknown | CI-NAU |
| DvRes-2 | *Pm21-F2* | Resistant | Unknown | GRA 961 | GBIS-IPK |
| DvRes-3 | *Pm21-F3* | Resistant | Italy | GRA 1114 | GBIS-IPK |
| DvSus-1 | *Pm21-NF1* | Susceptible | Greece | GRA 2738 | GBIS-IPK |
| DvSus-2 | *Pm21-NF2* | Susceptible | Unknown | GRA 962 | GBIS-IPK |
| DvSus-3 | *Pm21-NF2* | Susceptible | Italy | GRA 1105 | GBIS-IPK |
| DvSus-4 | *Pm21-NF3* | Susceptible | Former Soviet Union | PI 598390 | GRIN |
| DA6V#1 | *Pm21-NF3* | Susceptible | Developed by Sears | GSTR 315 | GRIN |
| DA6V#3 | *Pm21-NF2* | Susceptible | Developed by Lukaszewski | TA5618 | WGGRC-KSU |

<sup>a</sup> *Pm21-NF1* ~ *Pm21-NF3* were non-functional alleles of *Pm21* whose corresponding functional alleles were designated as *Pm21-F1* ~ *Pm21-F3*, respectively. In this study, *Pm21-F2* and *Pm21-F3* were successfully characterized, whereas *Pm21-F1* was not found in *D. villosum* accessions tested.

### Evaluation of powdery mildew resistance

All plants of *D. villosum* and wheat at one-leaf stage were inoculated with *Bgt* isolate YZ01, a predominant isolate collected from Yangzhou by dusting fungal conidiospores from susceptible wheat cv. Yangmai 9 onto the leaves (He et al. 2016), and powdery mildew responses were assessed as resistant or susceptible at eight days post-inoculation.

### Allelic test

The susceptible *D. villosum* line DvSus-1 was crossed with other susceptible ones (DvSus-2, DvSus-3 and DvSus-4), and three different F_1_ hybrids (DvSus-1/DvSus-2, DvSus-1/DvSus-3 and DvSus-1/DvSus-4) were obtained. The susceptible wheat-*D. villosum* addition line DA6V#1 was crossed with another susceptible addition line DA6V#3 to generate F_1_ hybrid DA6V#1/DA6V#3. Powdery mildew responses of F_1_ plants were assessed at eight days post-inoculation.

### Isolation of *Pm21* alleles

Genomic DNA was extracted from fresh leaves of seedlings by the CTAB method (Murray and Thompson 1980). PCR amplification was performed in Peltier thermal cycler (Bio-Rad, USA) in 50 μl volume containing 1× LA PCR buffer, 0.4 mM of each dNTP, 2 μM of each primer (forward primer: 5’-CTCACCCGTTGGACTTGGACT-3’; reverse primer: 5’-CTCTCTTCGTTACATAATGTAGTGCCT-3’), 1 unit of LA *Taq* DNA polymerase (TaKaRa, Japan), and 100 ng genomic DNA. PCR was carried out with an initial denaturation at 94°C for 3 min, 35 cycles of 20 s at 94°C, 30 s at 60°C, 3 min at 68°C, and a final extension for 5 min at 72°C. PCR products were cloned into the vector pMD18-T (TaKaRa, Japan), and then sequenced by the Sanger dideoxy sequencing method. To obtain exact sequences, each allele was isolated by three independent PCR, followed by sequencing. Full-length cDNA of each allele was further confirmed by sequencing of RT-PCR products. A phylogenetic tree was constructed by the neighbor-joining method in MEGA4 software (Tamura et al. 2007). Multiple sequence alignment analysis was performed by the CLUSTAL W tool (Thompson et al. 1994), and displayed using the GeneDoc tool (http://www.softpedia.com/get/Science-CAD/GeneDoc.shtml).

### Quantitative RT-PCR (qPCR) analysis

Total RNA was isolated from leaves of the seedlings with or without *Bgt* inoculation using TRIzol reagent (Life Technologies, USA). The first-strand cDNA was then synthesized from 2 μg of total RNA using a PrimeScript^™^ II 1st Strand cDNA Synthesis Kit (TaKaRa, Japan). qPCR was performed in an ABI 7300 Real Time PCR System (Life Technologies, USA) as previously described (He et al. 2018). A pair of universal primers (forward primer: 5’-TGAGTCTTCTAAACATCATTGC-3’; reverse primer: 5’-CACAACATGAACCTCGTCGT-3’) was used for detection of *Pm21* alleles. The wheat actin gene *TaACT* was used as the reference gene as reported (Bahrini et al. 2011). All reactions were run in three technical replicates for each cDNA sample.

## Results

### Powdery mildew responses of DvSus-1 ∼ DvSus-4, DA6V#1 and DA6V#3

Powdery mildew responses of four *D. villosum* lines (DvSus-1 ∼ DvSus-4) and two wheat-*D. villosum* addition lines (DA6V#1 and DA6V#3) were assessed at one-leaf stage with *Bgt* isolation YZ01. The results showed that all plants tested were susceptible (Fig. 1), which was in accordance with previous reports (He et al. 2017; Qi et al. 1998; Liu et al. 2011).

**Fig. 1.**
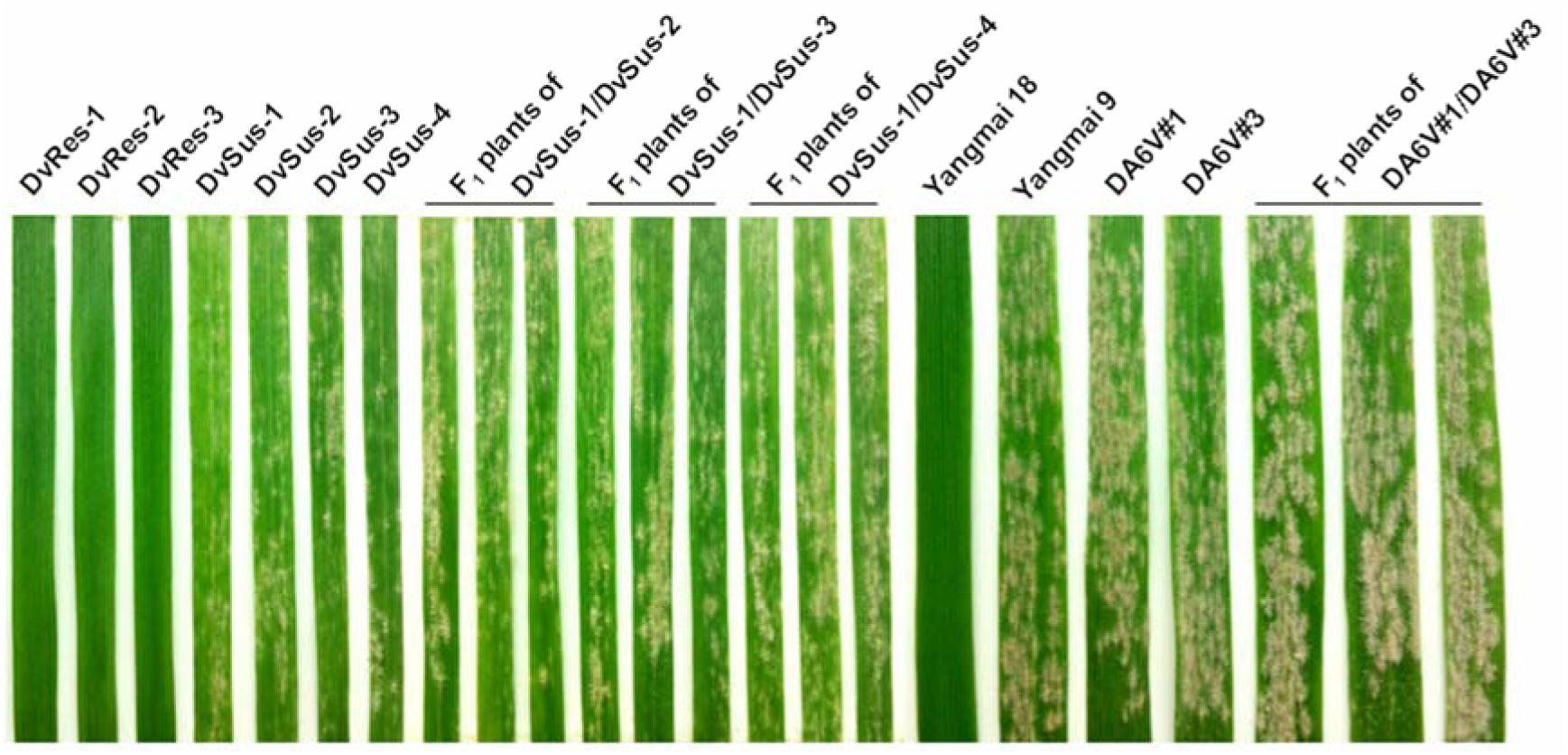
Powdery mildew responses of different *D. villosum* lines, wheat genetic stocks, and F_1_ plants of the crosses DvSus-1/DvSus-2, DvSus-1/DvSus-3, DvSus-1/DvSus-4, Dv6V#1/ Dv6V#3 to *Bgt* isolate YZ01 at one-leaf stage.

### Allelic test of susceptible factors in *D. villosum* and wheat background

To detect whether the susceptible factors are allelic, powdery mildew responses of F_1_ plants of four different crosses, including DvSus-1/DvSus-2, DvSus-1/DvSus-3, DvSus-1/DvSus-4, and DA6V#1/DA6V#3, were detected. The results demonstrated that all the F_1_ hybrids were susceptible to powdery mildew (Fig. 1), indicating no allelic complementation observed. Therefore, loss-of-function mutations in the susceptible *D. villosum* lines and wheat-*D. villosum* addition lines occurred in *Pm21* alleles.

### Quantitative RT-PCR (qPCR) analysis of *Pm21* alleles

To detect the relative transcription levels of *Pm21* alleles in different materials, qPCR was carried out. The data showed that *Pm21* alleles were all constitutively transcribed at relatively equal levels and displayed similar responses to *Bgt*, compared with the resistant *D. villosum* line DvRes-1 (Fig. 2). It was indicated that loss-of-functions of *Pm21* alleles might be caused by mutations in their encoding sequences rather than by the difference in their transcription patterns.

**Fig. 2.**
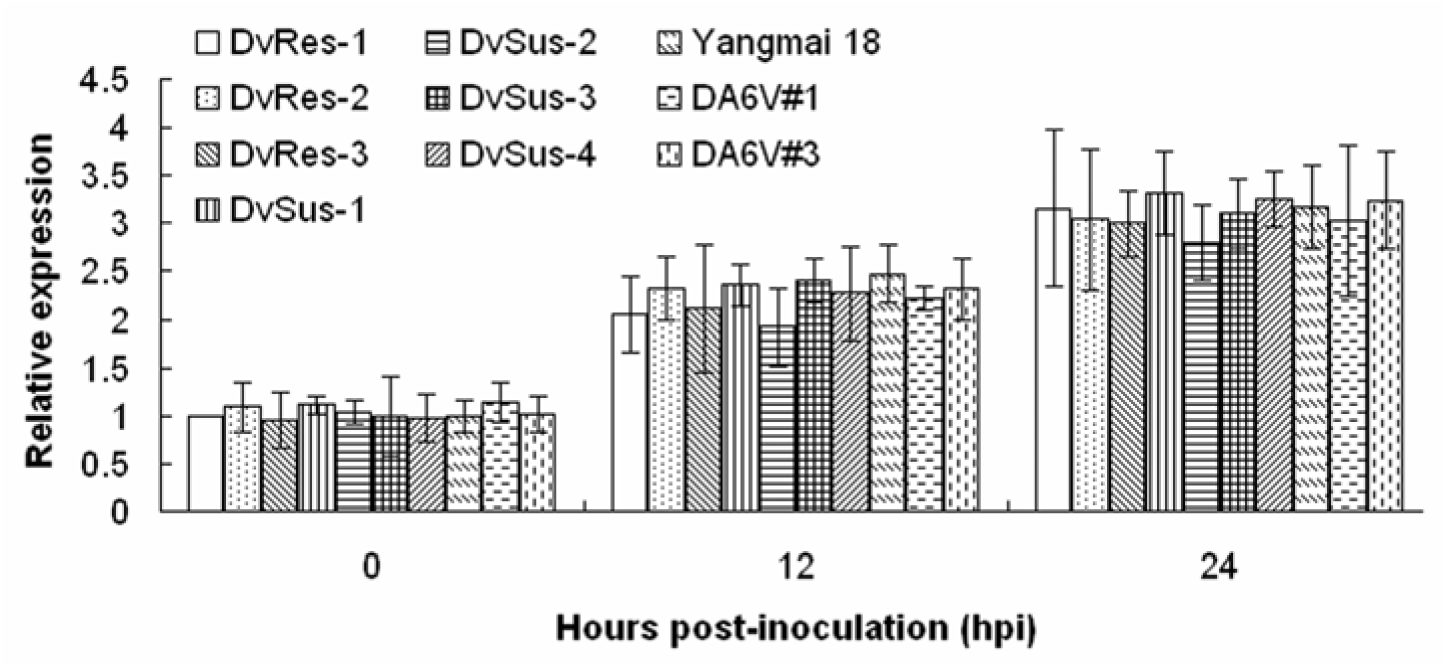
Transcription patterns of *Pm21* alleles in the leaves of the susceptible *D. villosum* lines and wheat genetic stocks inoculated with *Bgt* isolation YZ01, using quantitative RT-PCR (qPCR). The wheat *actin* (*TaACT*) gene was used as reference gene. The transcriptional values of *Pm21* allele in the *D. villosum* line DvRes-1 was set to 1 at 0 hpi. The error bars indicate standard deviation (SD) from three technical replicates.

### Sequence characteristics of non-functional *Pm21* alleles

To reveal the sequence characteristics, non-functional alleles of *Pm21* were isolated from the four susceptible *D. villosum* lines. The result showed that the genomic sequence of non-functional allele *Pm21-NF1* in DvSus-1 was 3,699 bp. The corresponding open reading frame (ORF) was 2,730 bp that had 98 single nucleotide polymorphisms (SNPs) contrasted to *Pm21*. However, *Pm21-NF1* only had two specific variation sites in contrast to 38 non-redundant alleles isolated from resistant *D. villosum* accessions reported recently (He et al. 2018). The first one was a G-to-T transversion at the position 61 that leaded to an amino acid change (A21S) in the CC domain. The second one was a A-to-G transition at the position 821 that caused another amino acid change (D274G), the second aspartate (D) in kinase-2 motif (consensus sequence: LLVLDDVW) of the NBS domain (Meyers et al. 1999) (Fig. 3A). Further analysis showed that the second D was conserved in all the 19 disease resistance proteins characterized from wheat, barley and *Arabidopsis* (Fig. 4).

**Fig. 3.**
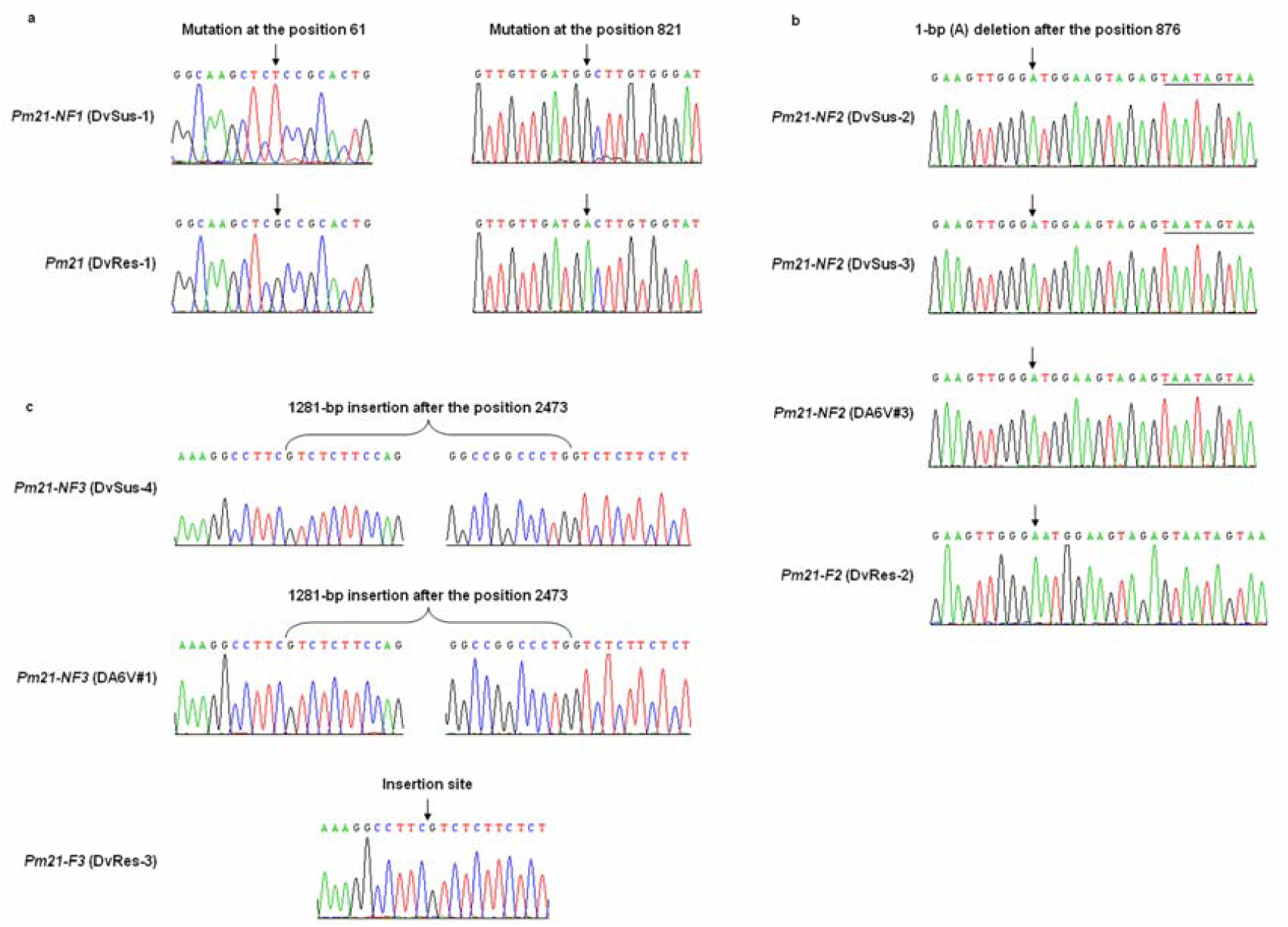
Detection of mutations in the non-functional *Pm21* alleles. **a** Mutations of *Pm21-NF1* in DvSus-1 contrasted to *Pm21* in DvRes-1. **b** Mutations of *Pm21-NF2* in DvSus-2, DvSus-3 and DA6V#3 contrasted to *Pm21-NF2* in DvRes-2. **c** Mutations of *Pm21-NF3* in DvSus-4 and DA6V#1 in contrast to *Pm21-F3* in DvRes-3. SNPs, tandem premature stop codons and insertion sequences are shown by *arrows*, underlines and brackets, respectively.

**Fig. 4.**
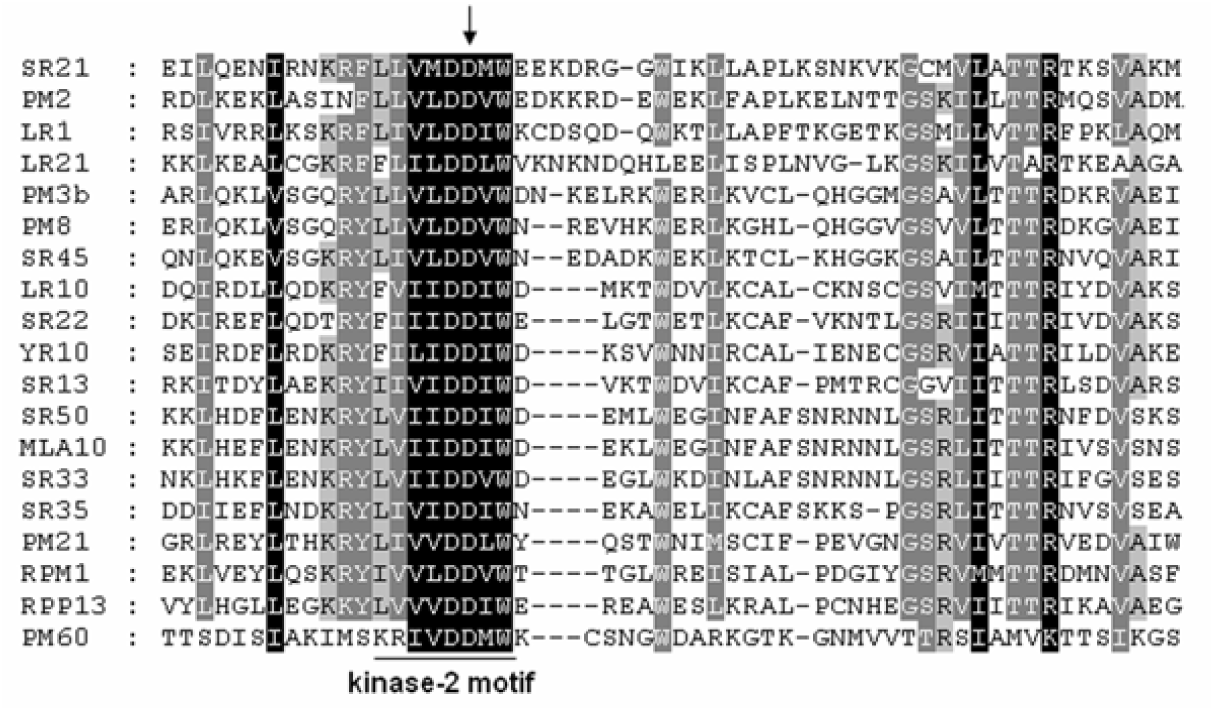
Multiple sequence alignment of the surrounding sequences of kinase-2 motif (hhhhDD) of plant disease resistance proteins. The second aspartate (D) of kinase-2 motif is pointed by an *arrow*

The genomic sequence of non-functional allele *Pm21-NF2* in DvSus-2 was 3,698 bp. The ORF of *Pm21-NF2* had a 1-bp (A) deletion after the position 876 that leaded to three tandem premature stop codons (TAA-TAG-TAA) in the NBS-encoding region and formed a truncated protein with 296 aa in length (Fig. 3B). The sequence of *Pm21* allele in DvSus-3 was identical to that in DvSus-2, which indicated the same origin.

The non-functional allele *Pm21-NF3* in the susceptible *D. villosum* line DvSus-4 were 4,988 bp in length. *Pm21-NF3* contained a 1281-bp insertion that is a kind of repeat sequence present in the genome of wheat cv. Chinese Spring (Fig. 3C). The insertion in *Pm21-NF3* introduced a premature stop codon, which leaded to a truncated LRR domain.

*Pm21* alleles in wheat-*D. villosum* addition lines DA6V#1 and DA6V#3 were also cloned and sequenced in this study. Interestingly, the non-functional *Pm21* alleles in DA6V#1 and DA6V#3 were identical to that in DvSus-4 and DvSus-2/DvSus-3, respectively (Fig. 3B and Fig. 3C). It was indicated that the non-functional *Pm21* alleles in the two wheat genetic stocks were both derived from their *D. villosum* donors.

### Molecular tracing of the origins of non-functional *Pm21* alleles

To trace the origins of non-functional *Pm21* alleles, a neighbor-joining phylogenetic tree was constructed based on the sequences of *Pm21* alleles (Fig. 5). The data showed that DvSus1, DvSus2, DvSus-3, DvRes-2 and GRA1164 were clustered in a clade. In contrast to the alleles in DvRes-2 and GRA1164, the non-functional allele *Pm21-NF1* in DvSus1 had eight and ten SNPs, and *Pm21-NF2* in DvSus2/DvSus-3 had one and three SNPs, respectively. It was indicated that the non-functional allele *Pm21-NF2* in DvSus2/DvSus-3 might originate from the corresponding functional allele *Pm21-F2* in DvRes-2 that was derived from the accession GRA961 (Fig. 6), whereas the origin of *Pm21-NF1* in DvSus1 could not be traced yet in the accessions tested.

**Fig. 5.**
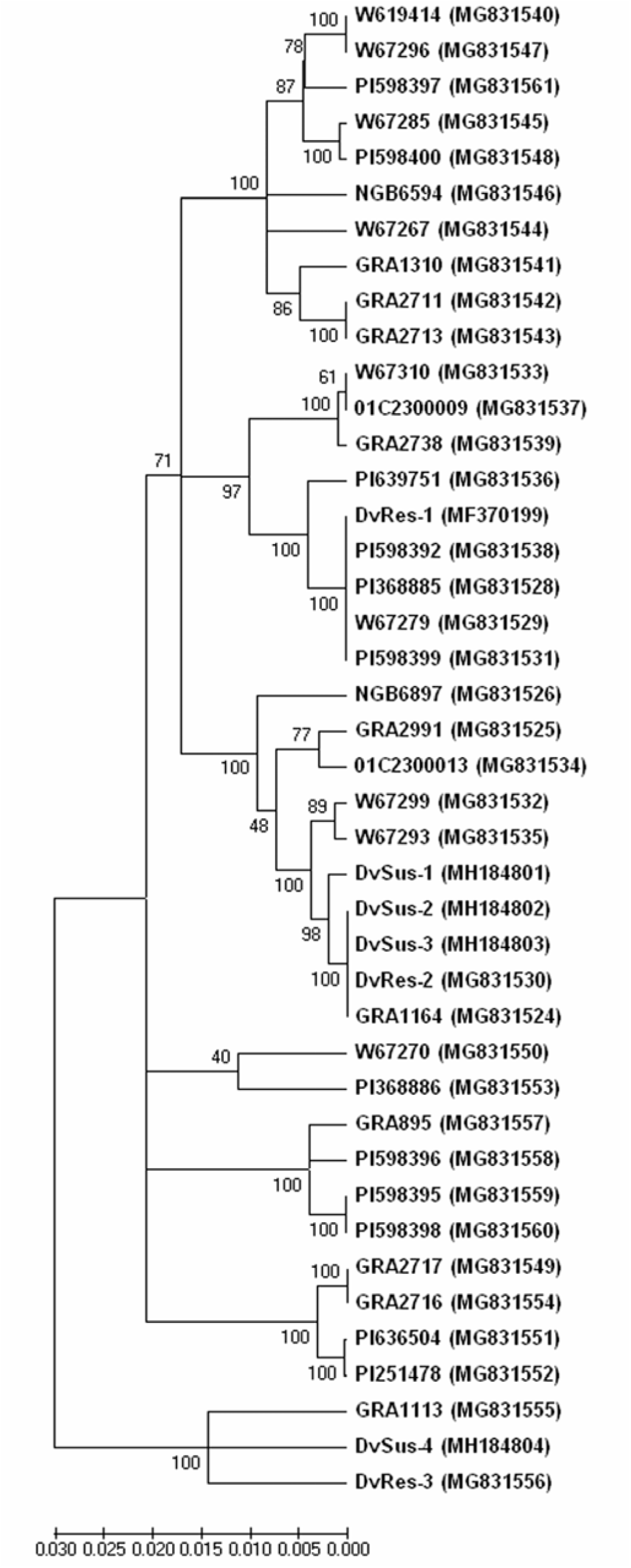
Phylogenetic tree based on the sequences of non-functional *Pm21* alleles obtained in this study, together with other *Pm21* alleles reported in our recent work (He et al. 2018). The accession numbers of alleles are shown in brackets.

**Fig. 6.**
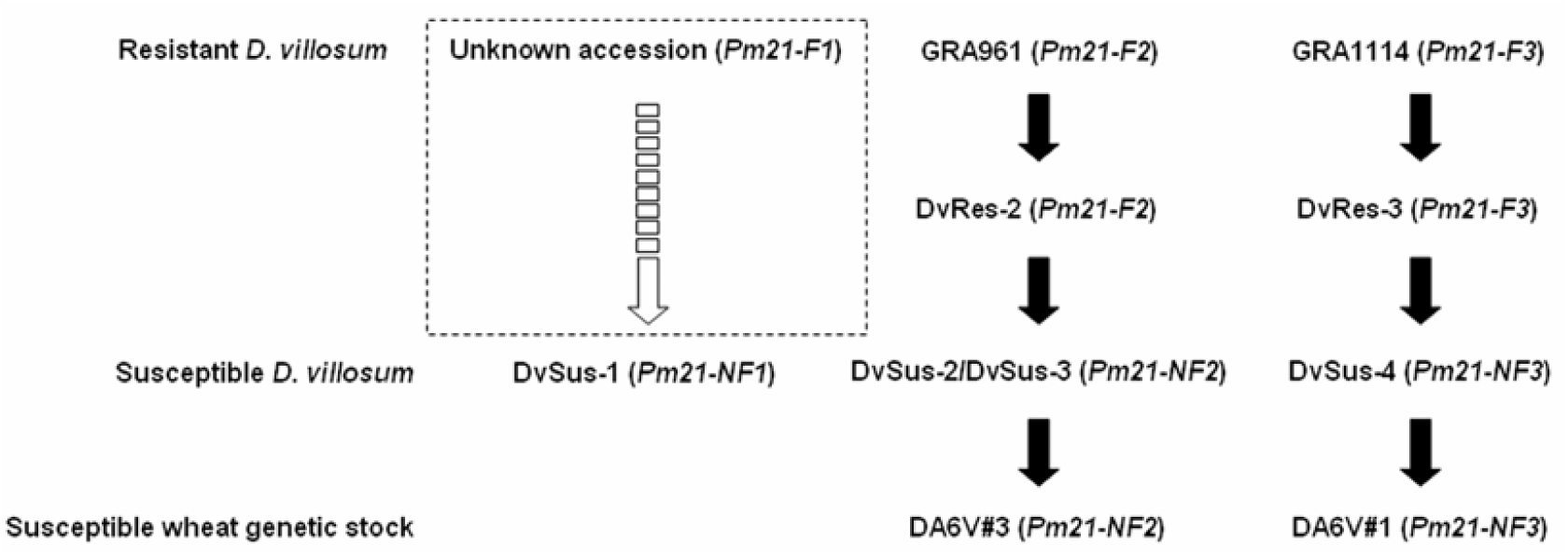
Origins of the non-functional alleles *Pm21-NF1* ∼ *Pm21-NF3*, corresponding to functional alleles *Pm21-F1* ∼ *Pm21-F3*, respectively. Among them, *Pm21-F1* was not detected in *D. villosum* accessions that is marked by a dashed box.

The phylogenetic tree also showed that DvSus-4, DvRes-3 and GRA1113 were clustered in an independent clade (Fig. 5). Sequence analysis demonstrated that, except the insertion sequence, *Pm21-NF2* in DvSus-4 had five SNPs contrasted to that in GRA1113, but no difference with that in DvRes-3. It was suggested that the non-functional allele *Pm21-NF3* in DvSus4 might originate from the corresponding functional allele *Pm21-F3* in DvRes-3 that was derived from the accession GRA1114 (Fig. 6).

## Discussion

*Pm21*, originating from *D. villosum*, confers highly effective resistance to wheat powdery mildew at the whole growth stages. To genetic map *Pm21*, 46 *D. villosum* accessions were investigated to find susceptible resource; however, none was obtained (Qi et al. 1998). Hence, since 1998, researchers believe that all *D. villosum* resources are resistant to powdery mildew. Recently, four seedling-susceptible *D. villosum* lines (DvSus-1 ∼ DvSus-4) showing complex resistance associated with growth stage were identified from different natural populations, which allows to fine map and clone *Pm21* (He et al. 2017; He et al. 2018). In this study, the characteristics of *Pm21* alleles in these susceptible lines were further investigated by Sanger sequencing. The data demonstrated that the variation of *Pm21* alleles in the four seedling-susceptible *D. villosum* lines involved in point mutation, deletion and insertion, respectively.

DvSus-2 and DvSus-3 contained the same allele *Pm21-NF2* that involved in a 1-bp deletion and leaded to a truncated PM21 protein. The length of the truncated protein (296 aa) is similar to that in the susceptible mutant lines Y18-S7 (281 aa) and Y18-S21 (309 aa) derived from EMS-induced wheat cv. Yangmai 18 carrying *Pm21*. In DvSus-4, the non-functional allele *Pm21-NF3* had a 1281-bp insertion of repeat sequence that leaded to lose the last four LRR motifs (13th to 16th). LRR domain of disease resistance protein plays an important role in perception of pathogen effector. Recent observation on three susceptible mutants of Yangmai 18 (Y18-S8, Y18-S30 and Y18-S47) showed that each mutation in the 14th LRR motifs, leading to premature stop codon or amino acid change, can impair the *Pm21* resistance (He et al. 2018). It was suggested that an important site(s) for perception of *Bgt* effector is lost in the truncated protein encoded by *Pm21-NF3*. Taken together, it is supported that both *Pm21-NF3* and *Pm21-NF3* were lost the resistance to powdery mildew.

Allelic tests on the crosses, DvSus-1/DvSus-2, DvSus-1/DvSus-3 and DvSus-1/DvSus-4, suggested that all mutations leading to lose resistance occurred in the alleles. Therefore, it was proposed that the allele *Pm21-NF1* in DvSus-1 was also non-functional. Contrasted to *Pm21, Pm21-NF1* had a number of SNPs; however, only two (G61T and A821G) were specific in all tested *Pm21* alleles that cause amino acid changes (A21S and D274G). A21S lied in the CC domain; however, whether it could impair the *Pm21* resistance remains unclear. D274 was the second aspartate (D) of kinase-2 motif (also called Walker B motif; consensus sequence: LLVLDDVW) in the NBS domain that is considered to act as the catalytic base for ATP hydrolysis and activation of disease resistance protein (Meyers et al. 1999; Tameling et al. 2006). Here, we investigated 19 plant disease resistance proteins and found that the second D of kinase-2 motif was highly conserved. This result supported that the amino acid change D274G might lead to loss-of-function of *Pm21-NF1*.

In this research, the origins of *Pm21-NF2* and *Pm21-NF3* were also successfully traced by sequence analysis. Except the mutation sites, *Pm21-NF2* and *Pm21-NF3* were identical to *Pm21-F2* and *Pm21-F3* isolated from the resistant *D. villosum* lines DvRes-2 and DvRes-3, derived from the accessions GRA961 and GRA1114, respectively. Therefore, it was suggested that *Pm21-F2* and *Pm21-F3* are the wild-type alleles corresponding to *Pm21-NF2* and *Pm21-NF3*, respectively.

Previously, researchers observed that wheat-*D. villosum* addition lines DA6V#1 and DA6V#3 were susceptible to powdery mildew (Qi et al. 1998; Liu et al. 2011). Due to the utilization of colchicine, an effectively chemical mutagen (Gilbert and Patterson 1965), in the incorporation of alien genome, it is difficult to explain whether the susceptibilities of DA6V#1 and DA6V#3 came from colchicine treatment or alien donors. In the present study, via allele sequencing, we revealed that *Pm21* alleles in DA6V#1 and DA6V#3 were identical to *Pm21-NF3* and *Pm21-NF2*, respectively. These results suggested that both of the variations of *Pm21* alleles in DA6V#1 and DA6V#3 originated from their *D. villosum* donors.

